# Nonparametric Anomaly Detection on Time Series of Graphs

**DOI:** 10.1101/2019.12.15.876730

**Authors:** Dorcas Ofori-Boateng, Yulia R. Gel, Ivor Cribben

## Abstract

Identifying change points and/or anomalies in dynamic network structures has become increasingly popular across various domains, from neuroscience to telecommunication to finance. One of the particular objectives of the anomaly detection task from the neuroscience perspective is the reconstruction of the dynamic manner of brain region interactions. However, most statistical methods for detecting anomalies have the following unrealistic limitation for brain studies and beyond: that is, network snapshots at different time points are assumed to be independent. To circumvent this limitation, we propose a distribution-free framework for anomaly detection in dynamic networks. First, we present each network snapshot of the data as a linear object and find its respective univariate characterization via local and global network topological summaries. Second, we adopt a change point detection method for (weakly) dependent time series based on efficient scores, and enhance the finite sample properties of change point method by approximating the asymptotic distribution of the test statistic using the sieve bootstrap. We apply our method to simulated and to real data, particularly, two functional magnetic resonance imaging (fMRI) data sets and the Enron communication graph. We find that our new method delivers impressively accurate and realistic results in terms of identifying locations of true change points compared to the results reported by competing approaches. The new method promises to offer a deeper insight into the large-scale characterizations and functional dynamics of the brain and, more generally, into intrinsic structure of complex dynamic networks.

## 1 Introduction

Identifying and analyzing change points and/or anomalies (which we use interchangeably) has become an increasingly active area of research in network sciences (Host-Madsen and Zhang, 2018; Messer et al., 2018). For example, in financial trading, a regime shift in the network of transactions is frequently linked to various (often upcoming) insolvencies such as bankruptcies, defaults, and recessions while a change in the network topology of cryptocurrency transactions may suggest a potential money laundering scheme (e.g., Elliott et al., 2014; Vandermarliere et al., 2015; Abay et al., 2019; Akcora et al., 2020). Similar to other biological networks (Wig et al., 2011), the idea of studying the brain as a dynamic functional network is helpful in understanding the complex network organization of the brain and can lead to profound clinical breakthroughs (Bassett and Bullmore, 2006; Rubinov and Sporns, 2010).

Most currently available statistical methods for anomaly detection in dynamic networks have the following limitation: network snapshots at different time points are assumed to be independent (e.g., Peel and Clauset, 2015; Akoglu and Faloutsos, 2010; Harshaw et al., 2016). This assumption appears to be unrealistic in many applications. For example, cryptocurrency entities and their interactions in a network of transactions evolve in time but obviously daily snapshots of the same network cannot be assumed to be independent. This dependence or autocorrelation effect is well documented in the time series literature since the 1960s (Zellner, 1962; Wolff et al., 1967). However, in many cases, this effect is often overlooked in practice, which leads to unreliable and false conclusions. Neglecting the dependence among the network snapshots at different time points leads to inflated false-positive rates of change points or anomalies, especially for small and moderate sample sizes.

To overcome this limitation, we propose a new distribution-free framework, named Network Evolution Detection Method (*NEDM*), for anomaly detection in dynamic networks and evolution network structures of high dimensional time series. The setup for the proposed methodology entails the following: With each network snapshot as a graph object, we find its unique univariate characterization, for example, mean degree, clustering coefficient, and clique number. As a result, a series of possibly very high dimensional network snapshots is transformed into a time series of scalars. Given the temporal dependence of network snapshots, it is infeasible to assume that the resulting time series of linear characteristics is independent. Next, we adopt a change point detection method (Gombay, 2008) for (weakly) dependent time series that is based on efficient scores. To enhance the finite sample properties of detected change points, we approximate the asymptotic distribution of the test statistic with a sieve bootstrap procedure (Kreiss, 1988, 1992; Bühlmann et al., 1997). We derive asymptotic properties for the *NEDM* statistic, validate its performance in respect to competing anomaly detection methods via synthetic and real data experiments. We illustrate the utility of the *NEDM* by applying it to two input data types: two multivariate functional magnetic resonance imaging (fMRI) time series data (Cribben et al., 2012, 2013; Cribben and Yu, 2017) and a dynamic network of the Enron email communication (Park et al., 2012; Priebe et al., 2005; Diesner et al., 2005). For the sake of brevity, we defer the analysis of one fMRI data set to the Supplementary Material (Appendix).

In application, we find that the *NEDM* has the potential to unveil the time-varying cognitive states of both controls and subjects with neuropsychiatric diseases such as Alzheimer’s, dementia, autism and schizophrenia in order to develop new understandings of these diseases. By applying the *NEDM*, we can consider whole brain dynamics, which promises to offer deeper insight into the large scale characterizations of functional architecture of the whole brain.

To sum up, the proposed *NEDM* has the following unique and significant attributes:

1. It is, to the best of our knowledge, the first paper to consider estimating change points in any graph summary statistic for the time-evolving network structure in a multivariate time series context.
2. It can consider thousands of time series and, in particular, the case where *P*, the number of time series is much greater than *T* (*P* >> *T*), the number of time points.
3. Unlike existing methods it is not limited by assuming that network snapshots at different time points are independent.
4. It enhances the finite sample properties of the change point method by approximating the asymptotic distribution of the test statistic using the sieve bootstrap.
5. Although it is inspired by and developed for brain connectivity studies, it pertains to a general setting and can also be used in a variety of situations where one wishes to study the evolution of a high dimensional network over time.

The remainder of the paper is organized as follows. We introduce the *NEDM* in Section 3. A simulation study that examines the finite sample performance of our method via sieve bootstrapping (Kreiss, 1988, 1992; Bühlmann et al., 1997) is also covered in Section 3. In Section 4, we describe the properties for building synthetic data and then provide background information on the fMRI and Enron dynamic network data sets in Section 5. The performance of the proposed algorithm, on both the synthetic and real world data, is detailed in Section 6. Conclusion and future work is provided in Section 7. Finally, proofs, supplementary material and processes are deferred to the Supplementary Material.

## 2 Related Work

There exists a vast body of studies on dynamic network models across various disciplines (see, Barabási and Albert (1999) and references therein). One method based on the minimum description length (MDL) principle and compression techniques (Sun et al., 2007) flattens the adjacency matrices into binary strings and uses compression cost to derive data specific features. Another procedure (DELTACON) proposed by Koutra et al. (2013) relies on the similarity measures between a pair of equal node networks. However, anomalous points reported by this procedure tend to suffer from limited interpretability due to their lack of statistical quantifiers (such as critical numbers or *p*-values).

The first comprehensive treatment of high dimensional time series factor models with multiple change points in their second-order structure has been put forward by Barigozzi et al. (2018). To detect changes in the covariance matrix of a multivariate time series, Aue et al. (2009) introduced a method using a nonparametric CUSUM type test, and Dette and Wied (2016) proposed a test where the dimension of the data is fixed. Furthermore, Cribben et al. (2012, 2013) put forward a method for detecting changes in the precision matrices (or undirected graph) from a multivariate time series. In turn, Cribben and Yu (2017) introduced a graph-based multiple change point method for changes in the community network structure between high dimensional time series, called Network Change Point Detection, that uses an eigen-space based statistic for testing the community structures changes in stochastic block model sequences. In addition, Barnett and Onnela (2016) developed a method for detecting change points in correlation networks that, unlike previous change point detection methods designed for time series data, requires no distributional assumptions.

## 3 Methodology

In this section, we describe the proposed contribution which tracks structural changes within the network structure of data sets. We use the terms change point detection and anomaly detection as well as the terms graphs and networks interchangeably. Table 1 in the Supplementary materials details the notation for the rest of the paper.

**Table 1:** Notation

| Symbol | Description |
| --- | --- |
| $G = (V, E)$ | graph/network; $V$ set of vertices, $E$ set of edges |
| $q; \omega$ | window length, threshold parameter |
| $\mathbf{D}$ | $T \times P$ multivariate time series; $T$ number of time points, $P$ number of time series |
| $\mathbf{F}$ | $q$-row fold of $\mathbf{D}$ |
| $\mathbf{R}; \mathbf{A}$ | correlation matrix; adjacency matrix |
| $CLQNUM$ | clique number |
| $MD$ | mean node degree |
| $ACC$ | average clustering coefficient |
| $APL$ | average path length coefficient |
| $BOM$ | Barnett & Onnela’s change detection method |
| $KVFM$ | Koutra et al.’s DELTACON method |
| $NEDM$ | network-evolution detection method |
| $minLCC$ | minimum local clustering coefficient |
| $maxBETW$ | maximum node betweenness centrality |

### 3.1 Input Data: Graphs and Multivariate Time Series

Our proposed methodology is applicable to two types of the input data: data that are originally in a form of a graph, and multivariate time series that are used to construct a graph, based on a certain similarity measure, e.g., correlation. Since one of our primary motivating applications is multivariate fMRI time series, below we describe in details how networks can be constructed from such data sets.

#### Networks from a multivariate time series

In many applications such as, for instance, neuroscience and finance, input data are multivariate time series, and the first step consists of constructing a graph structure based on a user-selected (dis)similarity measure. That is, suppose ***D*** is a *T* × *P* multivariate time series, where *T* and *P* are the number of time points and the number of time series, respectively. From the multivariate ***D***, we take a *q*-row sample, 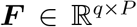, in a sequential “one-in, one-out” manner. This mode of data segmentation is known as the *overlapping/sliding window* technique (Keogh et al., 2001) and is ideal for maintaining the time dependency structure within ***D***, while taking as many samples as possible from the data matrix, ***D***. Because each *q*-row sample contains information from previous or successive samples, this segmentation procedure has the advantage of capturing all (and any) network structure disturbance. Note that, depending on the window length *q*, some data information may not be adequately captured because not every row in the data matrix has an equal number of appearances in all the ***F*** folds created. For instance, the first row (***D***_[1,]_) and the last row (***D***_[*T*,]_) are least likely to be included in the collection of all ***F***s. However, since we hypothesize that the changes points occur within the data and not at the end points (at the beginning and end of the time series) our detection procedure and results do not suffer.

Armed with ***F***, we compute a correlation matrix 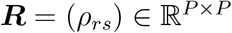 by quantifying the linear association between the *P* vertices in ***F***. With ***R*** and a pre-defined threshold *ω*^1^, we define the finite graph-associated adjacency matrix 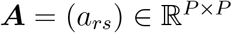 using

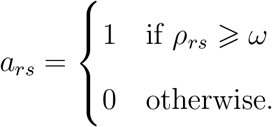

Given each adjacency matrix ***A***, we then construct a graph object *G_t_* = (*V_t_, E_t_*). (Here, to identify a minimum feasible number *P*, as a potential rule of thumb, a practitioner can employ approaches discussed, e.g., by Ozkaya et al. (2017).)

#### Input data as a graph structure

Alternatively, the original input data can take form of a graph *G_t_* observed at time point *t, t* = 1, 2,…. Such examples include communication networks (see, for instance, the Enron study in Sections 5 and 6), power grid networks, and the emerging blockchain technology. Indeed, one of the salient blockchain features is that all transactions are permanently recorded on distributed ledgers and publicly available. As a result, a blockchain graph *G_t_* can be constructed directly on each transaction, bypassing application of correlation and other similarity measures.

Finally, armed with the sequence of the graph objects *G_t_*, we then calculate various global and local graph summary statistics (Newman, 2003; Barabási and Pásfai, 2016). In particular, from each graph, we estimate the following graph summaries (*Y_t_*): Average Clustering Coefficient (*ACC*), Average Path Length (*APL*), Maximum Node Betweenness centrality (*maxBETW*), Clique number (*CLQNUM*), Mean Degree (*MD*) and Minimum Local Clustering Coefficient (*minLCC*).

### 3.2 Detection procedure and the sieve bootstrap

Following the dimension reduction and data structure simplification in Section 3.1, our next task is to identify the time(s) at which the regime shift(s) occur in the series of lower dimensional embeddings. In our case, we are interested in testing the following hypotheses:

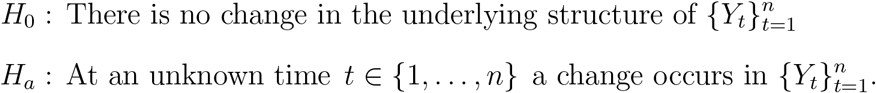

We assume that the appropriate model to fit to 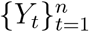 is a strictly stationary and purely non-deterministic autoregressive [*AR*(*p*)] model with Gaussian independent and identically distributed (i.i.d.) white noise *ε_t_*. The assumption of Gaussianity for *ε_t_* can be relaxed and substituted by the appropriate moment conditions (Gombay, 2008). (We run experiments on time series with non-Gaussian innovations and find that while the proposed change point detection is applicable to a non-Gaussian case, performance largely depends on deviations from the normality assumption and sample size. For instance, under the null hypothesis of no change, a nominal *α*-level of 0.05, and an AR(1) process with *φ* of 0.5, it takes approximately 100 observations to achieve an approximate size of the test of 0.05 for a case of *t*-distribution with 9 degrees of freedom; in turn, it requires about 200 observations from a uniform distribution to achieve a similar empirical size of the test.) In turn, an approximation of time series via AR(*p*) models, including the case of *p* → ∞, is widely used in theory and methodology of time series analysis (for overview see, for instance, Pourahmadi (2001); Shumway and Stoffer (2017), and references therein).

#### Remark

Note that under the considered problem of change point detection on networks, the network topological summary statistic 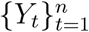 is estimated from the data. There currently exist no theoretical results on asymptotic properties of network statistics, especially in a conjunction with dynamic networks; that is, besides invoking a central limit theorem for network mean degree, nothing can be formally said on a linear process representation of 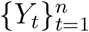 or its distributional properties. As a result, our approach is approximation-based and data-driven; that is, while we cannot provide theoretical bounds on the linear process approximation of 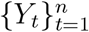, we can validate performance of the proposed *NEDM* against known ground truth change points.

In particular, we define *Y_t_* as

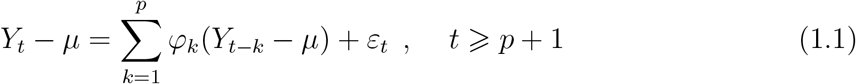

with **E**[*Y_t_*] = *μ* and let ***ξ*** = (*μ*, *σ*^2^, *φ*_1_,…, *φ_p_*)^*T*^. Given equation (1.1), we formally test the null hypothesis of no change

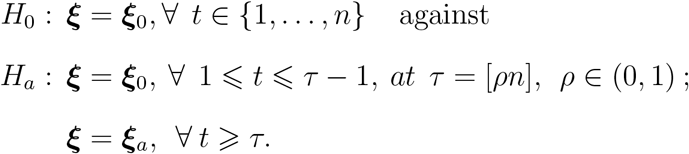

Most studies on change point detection calculate the pre-regime switch and post-regime switch parameter values of all possible change points *τ* ∈ (1, *n*), and then either use the strength of their differences to determine a regime switch or use these parameter values in the likelihood function (Picard, 1985; Inoue, 2001). However, we use a detection algorithm that involves a one-time parameter estimation and allows us to test for change in the individual elements of ***ξ***. In line with allowing for one-sided tests and for flexibility, we adopt the change point statistic of Gombay (2008) which utilizes the efficient score vector 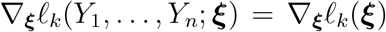 with *ℓ_k_* as the log-likelihood function on 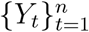. For for 1 ⩽ *r* ⩽ *p* + 2 denote 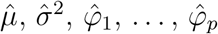 as the simultaneous solutions of the *p* + 2 equations *∂/∂ξ_r_ℓ_n_*(***ξ***) = 0, and let

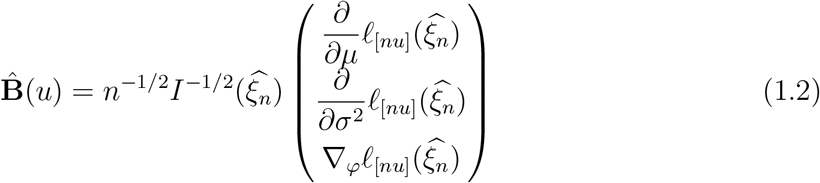

be a Gaussian process (a partial sums process approximation for the structure of ∇_***ξ***_ *ℓ_k_*(***ξ***)).

If *ξ_r_*, for 1 ⩽ *r* ⩽ *p* + 2, changes at *τ* = [*ρn*] with 0 < *ρ* < 1, then the estimator 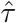 is defined by:

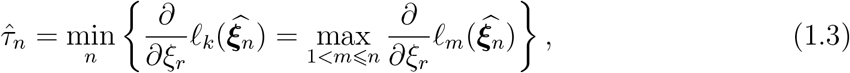

and

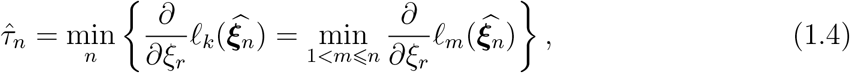

for one-sided tests (both left and right respectively); and for two-sided tests

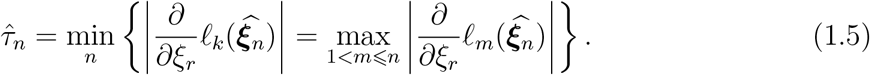

Proof for the consistency of 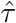 is provided in Gombay (2008). The asymptotic independence of the components of 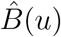 allows us define the change point test statistic for each *ξ_r_* ∈ ***ξ*** (see Gombay (2008) for the related discussion). Hence, we reject the null hypothesis, for *one-sided tests* along the sequence 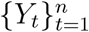, if

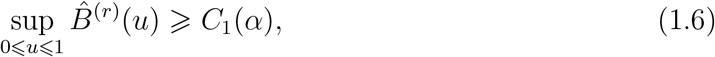

where *C*_1_(*α*) is calculated from

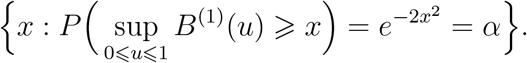

If we are interested in *two-sided* hypothesis testing, a change in *ξ_r_* (along the sequence 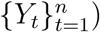 is acknowledged whenever

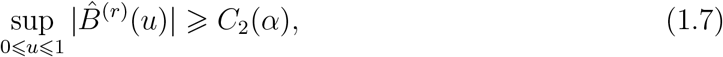

such that *C*_2_(*α*) is calculated from

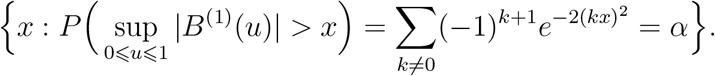

Convergence of the test statistic 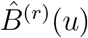 to its asymptotic distribution can be relatively slow. The Type I error estimates tend to be conservative with lower power of the test. Under the premise that 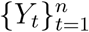 follows an *AR*(*p*) model with Gaussian i.i.d. white noise, we propose to adopt a sieve bootstrap procedure for constructing the distribution of the change point test statistic 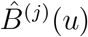 for finite samples. The idea of such a bootstrap for time dependent data – originally named AR(∞) bootstrap – goes back to the results of Kreiss (1988, 1992). The approach was later further investigated by Bühlmann et al. (1997) who coined the term *sieve* for this bootstrap method. The procedure is outlined in Algorithm 1, and its theoretical properties are stated in Theorem 1.

To compare finite sample performance of the asymptotic distribution to the sieve bootstrap method, we simulate data (with a total of 5000 Monte Carlo iterations) from an AR(1) model in equation 1.1 and evaluate the following: the size and power of the test [for precise details on the simulation procedure used, please see the Appendix]. The choice of the model coefficient (*φ*) depends on ensuring the assumption of weak stationarity for the simulated AR(1) series. Indeed note that as *φ* approaches 1, the time series gets closer to a random walk process, that is, we approach to a boundary case of violating the assumptions of weak stationarity, outlined for the change point statistic based on the efficient scores (Theorem 1 of Gombay, 2008). As a result, the performance of the change point detection method deteriorates.

Figure 1 depicts the size of the test for the simulated data. It shows that the asymptotic distribution behaves more conservatively compared to our sieve bootstrap distribution. Additionally, we find that the size is more conservative for the asymptotic distribution when *φ* is closer to +1 (*i.e*., the size is worse when *φ* = 0.9 compared to *φ* = 0.5). However for negative *φ* cases, we find that the size of the test under the asymptotic distribution approaches the declared 5% level of significance as *φ* approaches −1. Apart from the fact that the sieve bootstrap’s size is closer to the nominal rate than the asymptotic distribution, we find that as *φ* approaches +1 the size of the test steadily hovers around the declared 5% level of significance. This supports our assertion that the asymptotic distribution of the change point test statistic has relatively conservative Type I error, and validates our sieve bootstrap procedure for finite samples.

**Figure 1:**
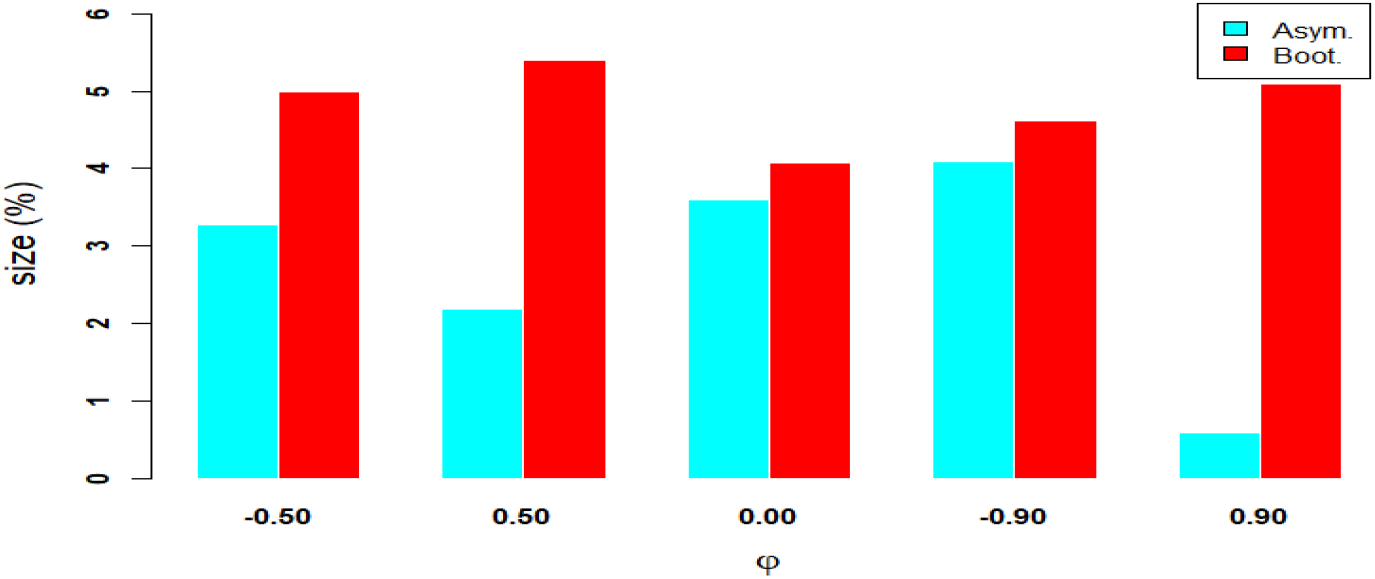
The size of the test plotted against *φ* for the asymptotic distribution and sieve bootstrap procedure when testing for a change in the mean of an *AR*(1) model using *T* = 100.

The results for power is displayed in Figure 2 and from this we notice that as *μ* increases, the power of the test also increases under both the asymptotic and the bootstrap distributions. In addition, as *φ* approaches +1, there is a drop in the power values for both the asymptotic and sieve bootstrap (with larger power values reported by our bootstrap procedure). Next to this, we find that as *φ* approaches −1 the power of the test improves for both the bootstrap and the asymptotic distribution; with the bootstrap again outperforming the asymptotic distribution.^2^

**Figure 2:**
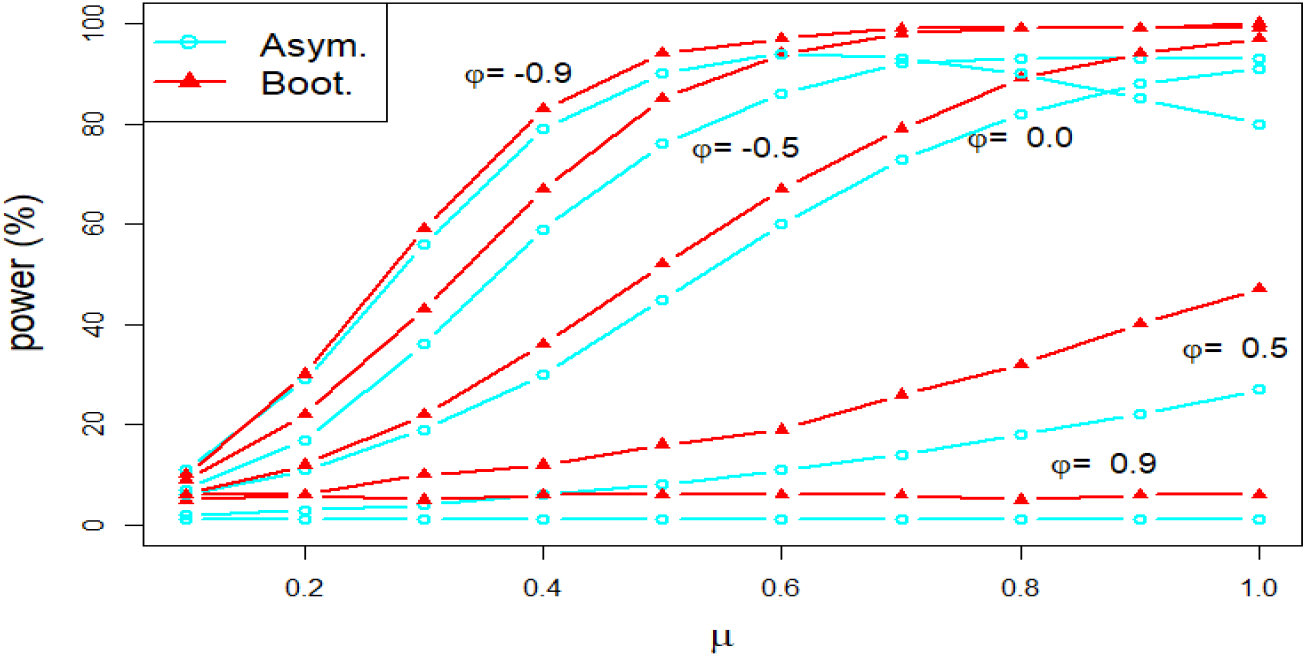
Power of the test plotted against *φ* for the asymptotic distribution and sieve bootstrap procedure when testing for a change in the mean of an *AR*(1) model with (*T* = 100).

##### Algorithm 1: Nonparametric sieve bootstrap procedure for the change point statistic under an *AR*(*p*) process

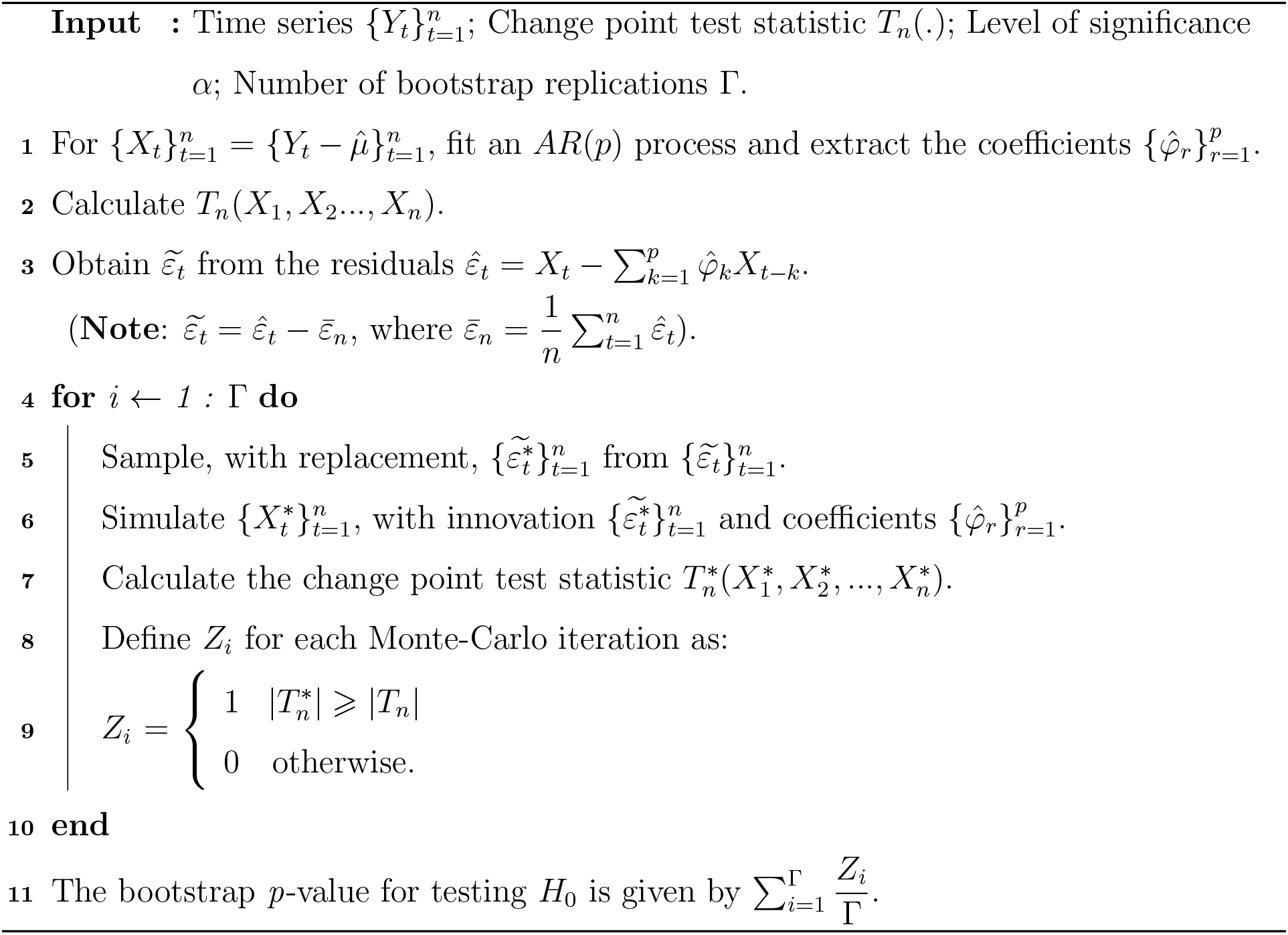

##### Theorem 1 (Sieve bootstrap)

*Let Y_t_ be an autoregressive process as defined in equation (1.1)* [*with* 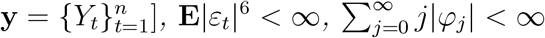, *and p*(*n*) = *o*((*n*/*log*(*n*))^1/4^). *With* 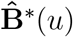 *as the bootstrap estimate for the p* + *2-dimensional Gaussian process* **B**(*u*), *and as n* → ∞, *we have*

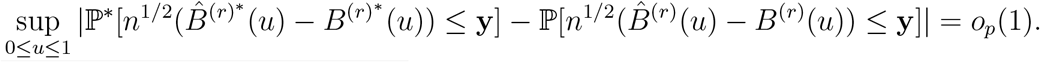

Proof of Theorem 1 is in the Appendix.

Suppose 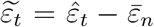, with 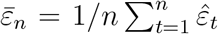, then the nonparametric sieve bootstrap estimate 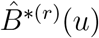 can be replaced with a hybrid parametric bootstrap estimate 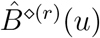 by generating a Gaussian sample 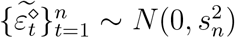, with 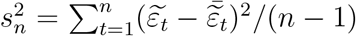 as the sample variance of 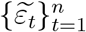. This yields a finite sample performance similar to that of 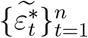, with reduced computing time.

##### Corollary 1

If *Y_t_* satisfies equation (1.1), then under *H*_0_

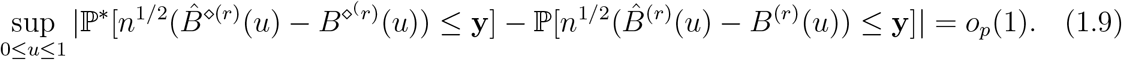

Proof of Corollary 1 is in the Appendix.

## 4 Simulations

To validate our *NEDM*, we compare it to the *BOM* (Barnett and Onnela, 2016) using simulated data. In particular, we provide a sensitivity analysis for the *NEDM* under various *q, ω* and ROI choices. We do not compare the *NEDM* to the *KVFM* (Koutra et al., 2013) because it does not provide a quantifier for the statistical significance (ac *p*-value) for the change point detected. Such a quantifier is necessary for deducing the power and size of the test for the methods under study. The simulation study covers two scenarios: a no-change point and a one-change point. As evaluation metrics, we use the true detection rates (i.e., power of the test) and the false alarm levels under a pre-defined significance level (i.e., size of the test). A total of 100 Monte Carlo simulations is carried out in each scenario under various window length *q* = {5, 10, 15, 20, 25, 30}, threshold parameter *ω* = {0.05, 0.1,0.15}, two different time series lengths (*T* = 200, 300) and for two graph node sizes (*P* = 5, 10). In addition, we provide similar analyses with *T* = 300, *P* = 50 in Appendix B of the Supplementary material.

The first simulation illustrates the baseline (no change point) scenario using a vector autoregression (VAR; Zellner, 1962; Hamilton, 1995) model; the VAR model is a generalization of the univariate AR process with more than one time-evolving component. Given the (*p* × 1) vector of time series variables ***F***_*t*_ = (*f*_1*t*_, *f*_2*t*_,…, *f_pt_*)^*T*^, the *w*-lag vector autoregressive (VAR(w)) process is defined as

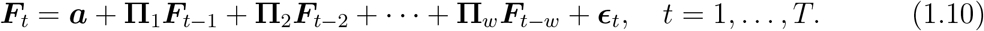

where **Π**_*i*_ is an (*p* × *p*) coefficient matrix and ***ϵ***_*t*_ is an (*p* × 1) unobservable mean white noise vector process with time invariant covariance matrix **Σ** (Zivot and Wang, 2007). The VAR model is used to reconstruct the linear inter-dependency element prevalent among multivariate time series applications such as fMRI data.

The second simulation, which depicts a one-change point scenario, is created by concatenating two data streams from distinct multivariate Gaussian distributions: (**D**_1_, **D**_2_) where **D**_1_ ~ *N*(***μ*** = 0, **Σ_1_** = (Σ_1*ij*_)) and **D_2_** ~ *N*(***μ*** = **0**, **Σ_2_** = (Σ_2*ij*_)) with

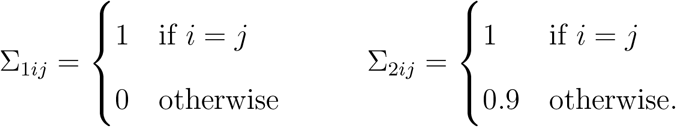

The results for the simulations are presented in Figures 4 and 5, and a discussion of the results is provided in Section 6.

## 5 Experimental Data

Next, we demonstrate the *NEDM*’s application to two input data types: multivariate fMRI time series and a portion of the Enron emails network. In the fMRI case study, we compare the performance of the *NEDM* against two other techniques for anomaly (change point) detection in the multivariate setting: the *KVFM* (Koutra et al., 2013) and the *BOM* (Barnett and Onnela, 2016). With the Enron networks, we only implement the *NEDM* with the *APL* network summary (because from our experiments the *APL*-based analysis produced the best outcome), and compare the change points we find with various events that characterized the timeframe of the Enron scandal. In addition, we compare the performance of the *NEDM* to results obtained by the *KVFM*. We do not compare our results with the *BOM* because the *BOM* is only applicable to multivariate time series data.

### 5.1 Case study: Anxiety fMRI data

The data was taken from an anxiety-inducing experiment (Cribben et al., 2012, 2013). The task was a variant of a well-studied laboratory paradigm for eliciting social threat, in which participants must give a speech under evaluative pressure. The design was an off-on-off design, with an anxiety-provoking speech preparation task occurring between lower anxiety resting periods. Participants were informed that they were to be given 2 *min* to prepare a 7 *min* speech, and that the topic would be revealed to them during scanning. They were told that after the scanning session they would deliver the speech to a panel of expert judges, though there was “a small chance” they would be randomly selected not to give the speech. After the start of fMRI acquisition, participants viewed a fixation cross for 2 *min* (resting baseline). At the end of this period, participants viewed an instruction slide for 15 *s* that described the speech topic, which was “why you are a good friend”. The slide instructed participants to be sure to prepare enough for the entire 7 *min* period. After 2 *min* of silent preparation, another instruction screen appeared (a relief instruction, 15 *s* duration) that informed participants that they would not have to give the speech. An additional 2 *min* period of resting baseline completed the functional run. Data were acquired and preprocessed as described in previous work (Wager et al., 2009). During the course of the experiment a series of 215 images were acquired (*TR* =2 *s*). In order to create ROIs, time series were averaged across the entire region. The data consists of 4 ROIs and heart rate for *n* = 23 subjects. The regions in the data were chosen because they showed a significant relationship to heart rate in an independent data set. The temporal resolution of the heart rate was 1 *s* compared to 2 *s* for the fMRI data. Hence, the heart rate was down-sampled by taking every other measurement.

### 5.2 Case study: Enron email networks

The Enron emails data set is a benchmark data set applied in numerous instances of anomaly detection (e.g., Peel and Clauset, 2015; Park et al., 2012; Priebe et al., 2005; Diesner et al., 2005). More information on this data set can be found online (http://www.cs.cmu.edu/~enron/). We used the cleaned version of the employee-to-employee email (sent and received) network over the period November 1998 to July 2001. We initialize each employee as a single node and aggregate the data by month. This implies that if there is at least one email between two employees within the month under study, an edge is connected to the respective nodes. Figure 3 displays the cumulative nature of the Enron network between November 1998 to July 2001, and the state of the network after two specific month/year periods. In total, we obtain 33 networks with 102 nodes in each network.

**Figure 3:**
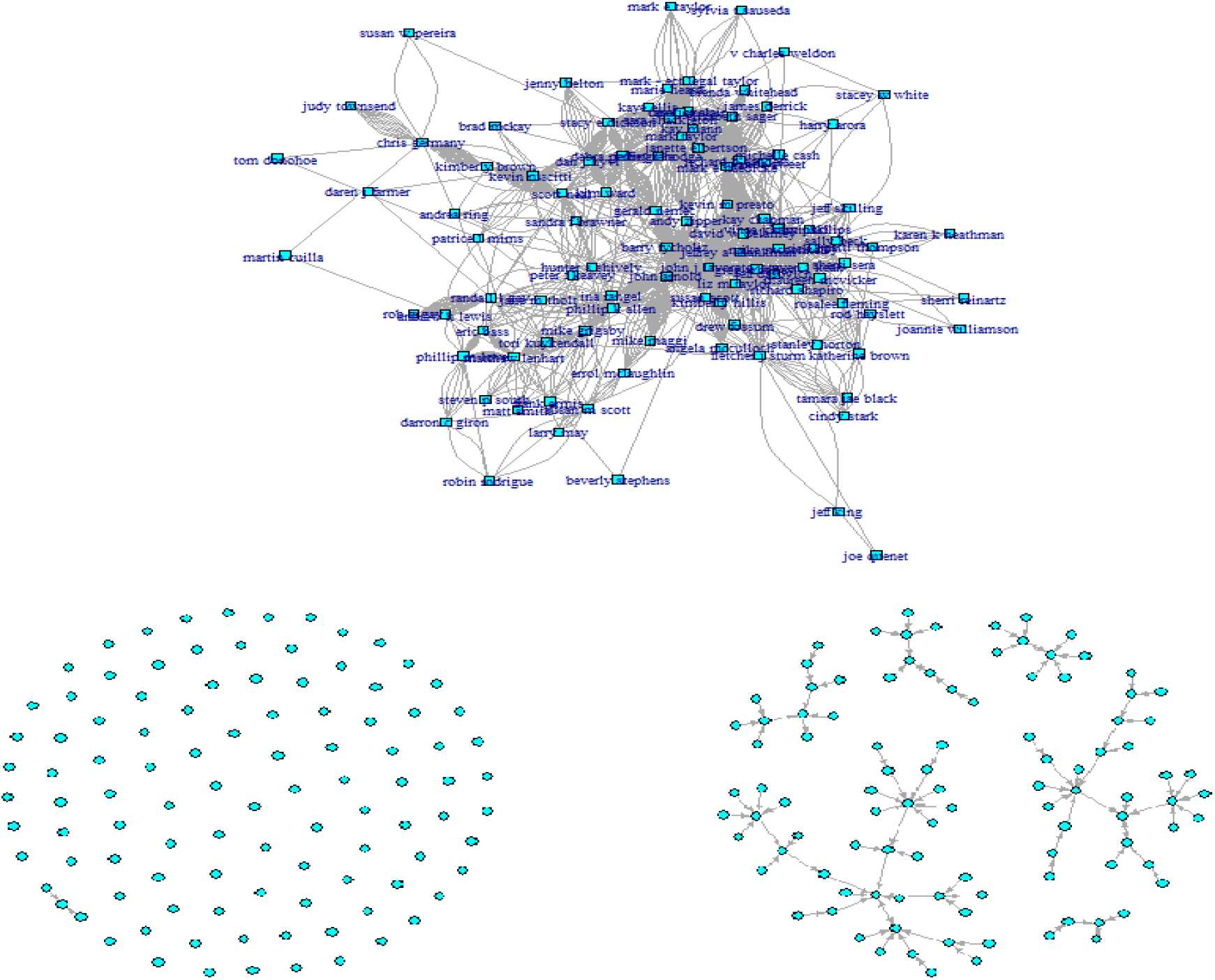
Enron employee-to-employee email network (November 1998 - July 2001). Top: Cumulative network from November 1998 to July 2001; Bottom (Left): Email network at November 1998, (Right): Email network at July 2001.

## 6 Results

The *NEDM* uses the following summary statistics: Average clustering coefficient (*ACC*), Average Path Length (APL), Maximum Node Betweenness Centrality (*maxBETW*), Clique number (*CLQNUM*), Mean degree (MD) and Minimum Local Clustering Coefficient (*minLCC*). As we already mentioned in the previous section, we compare our method to Barnett’s method (*BOM*) in the simulation study. However, we include results from *KVFM* for the Enron e-mail data set.

### 6.1 Simulation study

Figure 4 presents the results for the no-change point scenario. We find that as *q* increases the size of the test reported by all network summaries under the *NEDM* increase, leading to more liberal results. Moreover, we find that as *ω* increases, the size reported by all network summaries under the *NEDM* improve (and become closer to the 5% level of significance). In particular, we find that the size of the test reported by the *NEDM* under the *maxBETW* network summary almost always outperforms the size reported by the *BOM* (except for one situation when *w* = 0.05, *T* = 200 and *q* = 30). Furthermore, we notice that two other network summaries (*ACC* and *MD*) are highly sensitive to increasing *ω*, and that their size values improve substantially as *ω* increases from 0.05 to 0.15. For the *BOM* results, we find that the size of the test tends to be more liberal as *P* increases (the size increases from 18% to 20%), and that its performance is inferior to the *NEDM* with the *maxBETW* network summary. In conclusion, many of the graph summary statistics appear superior to the *BOM* in terms of the size of the test.

**Figure 4:**
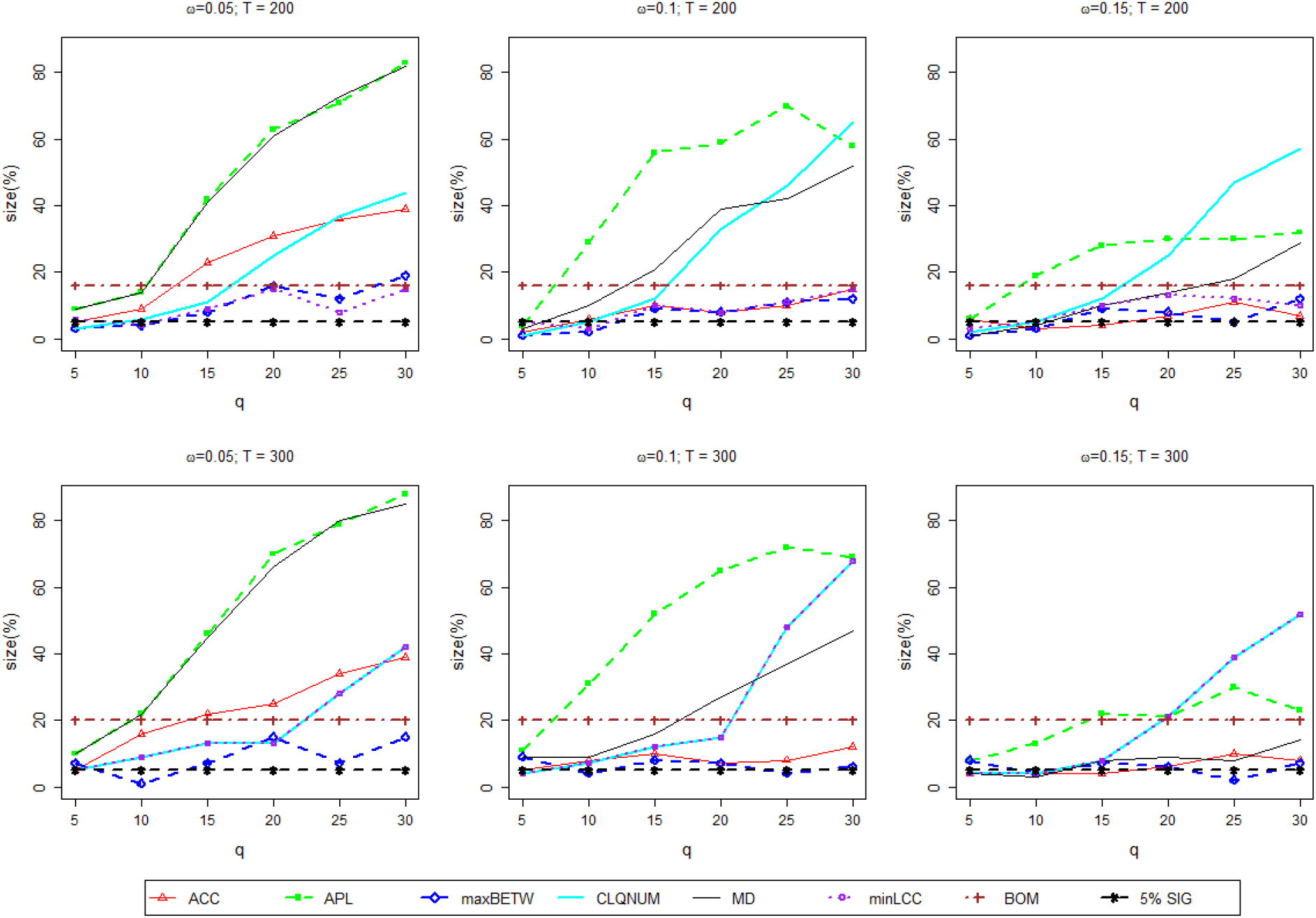
The size of the test for the no-change (change in mean) scenario with *T* = {200, 300}, *ω* = {0.05, 0.1, 0.15}, *q* = {5,10,15, 20, 25, 30} and *P* = 5.

The results for the one-change point simulation is presented in Figure 5. Overall, the *BOM* appears to attain the highest power (*i.e*., it correctly flags the one-change scenario every time). However, this performance is equally matched by the *NEDM* with the *maxBETW* network summary. In addition, for other network summaries utilized in the *NEDM*, we found that power drops as *ω* and *T* increase, and that power increases as *q* increases. Furthermore, we notice that as *T* increases from 200 to 300, the power reported by our *NEDM* with the *CLQNUM* summary statistic experiences a substantial drop, but this also improves as *q* rises.

**Figure 5:**
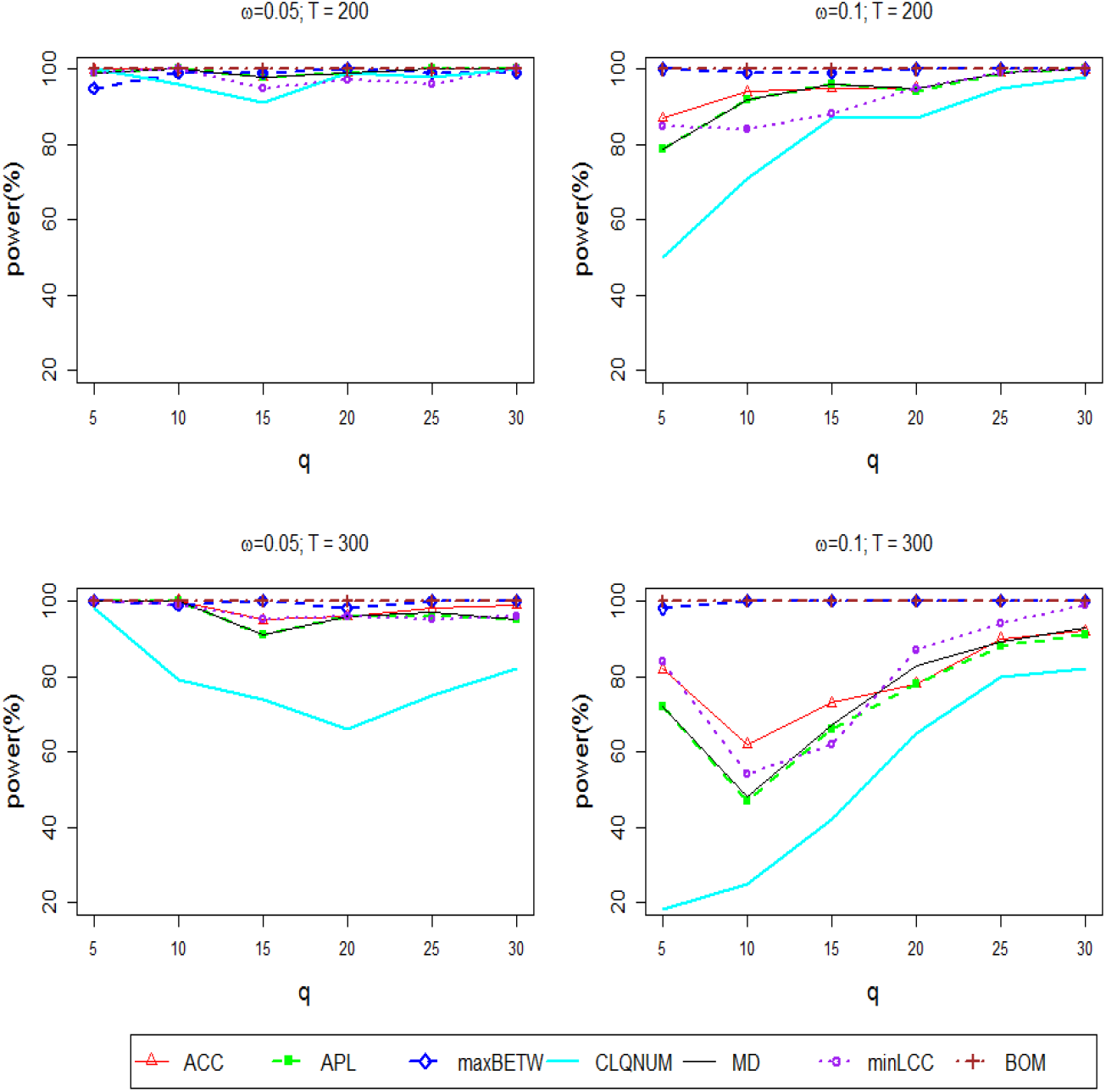
The power of the test for the one-change (change in variance) scenario with *ω* = {0.05, 0.1}, *q* = {5, 10, 15, 20, 25, 30}, *T* = {200, 300}, *τ* = {88, 156} and *P* = 10.

In summary, we found that both the power and the size of the test (detection ability) are sensitive to the choice of parameters (*q, ω* and *T*). In addition, while the *BOM* has excellent power, it suffers from liberal Type I errors. Overall, we found that the *NEDM* in combination with the *maxBETW* has the best performance in terms of maintaining the size of the test while also having excellent power.

### 6.2 Case study: Anxiety fMRI data

From our analyses, we obtained the results displayed in Figure 6 for the *NEDM* (with *ACC*, APL, *maxBETW* and *minLCC* as summary statistics), the *BOM* and the *KVFM*.

**Figure 6:**
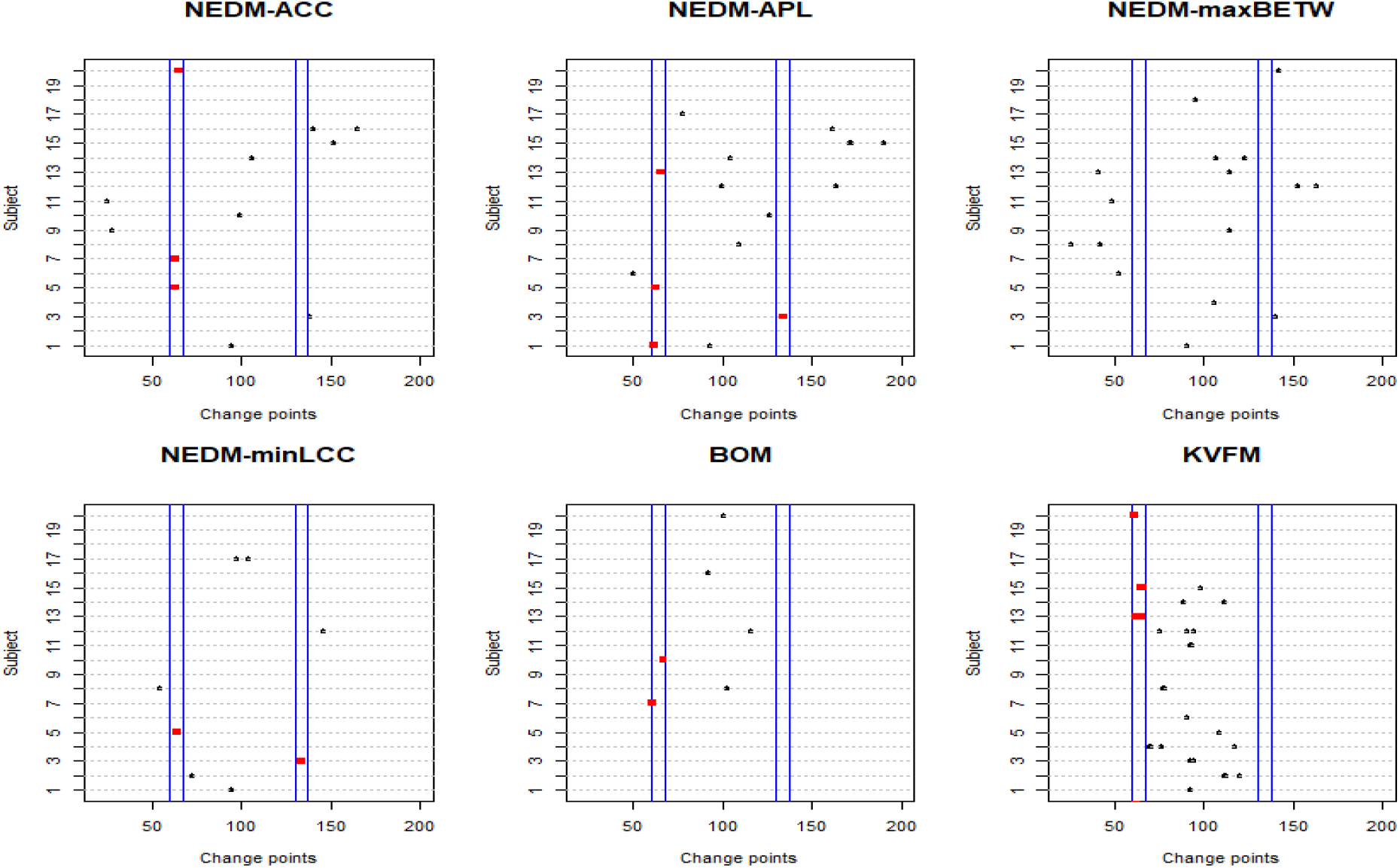
Change point detection for the anxiety fMRI data.

Results generated by the *NEDM* with *CLQNUM* and *MD* as the network topological summary statistic are deferred to the Supplementary material. Previous analyses of these data found change points at the times of the speech instruction slides, primarily time points 60 and 130 (Cribben et al., 2012, 2013). Additionally, all results obtained by the *NEDM* (plus the plot in the Supplementary material) found many of the expected change points at (and between) the time points of the speech instruction slides, and also during the speech preparation phase (time points 60-130).

While the *KVFM* found more changes points (as expected given it does not provide a quanitifier for statistical significance) at the time points of the speech instruction slides and also during the speech preparation phase, many of the change points for the subjects were very close to one another which makes them unrealistic for fMRI data. While the *BOM* also found some significant changes points at the speech instruction slides time points, the *NEDM* found significantly more. In addition, having change points so close to one another (as is the *KVFM* cases) makes it impossible to estimate the network structure between each pair of significant change points and also breaks down the ability to interpret and apportion the change phenomenon to a biological process (as in the case of the fMRI data used here). In turn, the framework of the *NEDM* allows for an estimation which depends on a visual display of the underlying dynamic brain networks with the advantage of noticing structural changes within these embeddings.

### 6.3 Case study: Enron email networks

We now turn to the anomaly detection problem for data in the form of a graph, that is, the Enron email network. From our list of network summary statistics, we utilize the *APL* because *APL* appears to be fairly sensitive to intrinsic properties of data in the form of dynamic networks. We link the results (Figure 7) obtained in this analysis to various events in the Enron scandal timeline. With the *NEDM*, the times for significant detection points occur at months 2, 7, 12, 15, 18 and 24. In comparison, the time period for month 2 (November 1998 to December 1998) is linked to the hire of Andrew Fastow as the finance chief. The time period of month 7 was from April 1999 to June 1999. This is around the time when Enron’s CFO was exempted by the Board of Directors from the company’s code of ethics so that he could run the private equity fund LJM1, and also around the time when the head of Enron’s West Coast Trading Desk in Portland Oregon, Timothy Biden, began his first experiment to exploit the new rules of California’s deregulated energy market.

**Figure 7:**
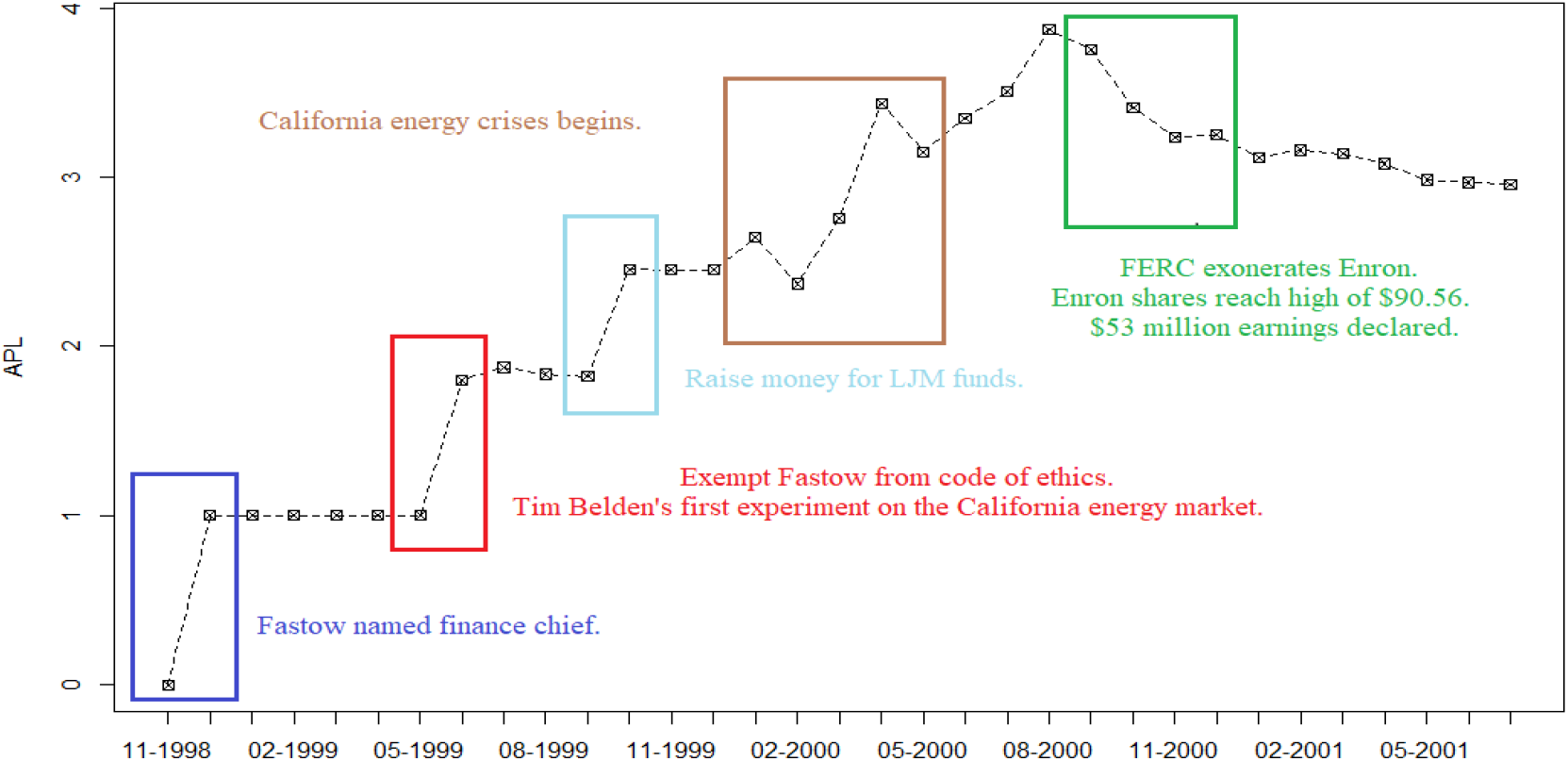
Detection of Enron Email Networks from November 1998 - July 2001.

Next to this, month 12 is linked to the period August 1999 to October 1999 which was around the time when Enron’s CFO started to raise money for two LJM funds (LJM1 and LJM2), which was later used to buy Enron’s poorly performing assets in order to make its financial statement look better. The time period for months 15 and 18 was from December 1999 to March 2000. This was close in time that the energy prices in California rose significantly and the power reserves became low, which was followed by the blackouts in metropolitan areas. Many believed that one of the reasons for California’s energy crisis was Enron’s trading, which leaded to the investigation of Federal Energy Regulatory Commission (FERC). This investigation was connected with our next detection point at month 24, which was from August 2000 to December 2000, since it was the time the FERC investigation exonerated Enron from wrongdoings in the California energy crisis. The time period around month 24 is a very interesting one not only because it is connected with FERC investigation, but also because Enron’s share price hit all-time high of $90.56 and then Enron used “aggressive” accounting to declare $53 million in earnings on a collapsing deal that hadn’t earned anything at all in profit.

Figure 2 in the Appendix displays the outcome of the *KVFM* analysis. Note that, because the *KVFM* is based on graph similarity scores, we only have results from December 1998 to June 2001 (Time point 0 is November 1998, and there are no more networks for comparison after July 2001). With the *KVFM*, anomalies are reported at times 2, 6 and 23. These times coincide with time periods [December 1998 - January 1999], [April 1999 - June 1999] and [September 2000 - November 2000]. In Figure 2 (in the Supplementary Materials), we notice that the *KVFM* is only able to flag 3 out 5 anomalous time points that the *NEDM* reported.

## 7 Conclusion

In this paper, we develop a new approach, the *NEDM*, for analyzing and modeling the network structure between (possibly) high dimensional multivariate time series from an fMRI study which consists of realizations of complex and dynamic brain processes. The method adds to the literature by improving understanding of the brain processes measured using fMRI. The *NEDM* is, to the best of our knowledge, the first paper to consider estimating change points for time evolving graph summary statistics in a multivariate time series context. Although this paper is inspired by and developed for brain connectivity studies, our proposed method is applicable to more general settings and can also be used in a variety of situations where one wishes to study the evolution of a high dimensional graph over time, e.g., in conjunction with telecommunication, financial (Cribben, 2019), and blockchain networks.

There are several novel aspects of the *NEDM*. First, it allows for estimation of graph summary statistics in a (possible) very high-dimensional multivariate time series setting, in particular, in situations where the number of time series is much greater than the number of time points (*P* >> *T*). Hence, in a biomedical neuroimaging setting, it can consider the dynamics of the whole brain or a very large number of brain time series, thereby providing deeper insights into the large-scale functional architecture of the brain and the complex processes within. Second, the *NEDM* is, to the best of our knowledge, the first piece of work to consider estimating change points of time evolving graph summary structure in a multivariate time series context. We introduced a novel statistical test for the candidate change points using the sieve bootstrap and showed that it outperformed the asymptotic distribution. However, as the *NEDM* is based on binary segmentation it is restricted by the minimum distance between change points.

It has been shown that neurological disorders disrupt the connectivity pattern or structural properties of the brain. Future work entails applying the *NEDM* to resting state fMRI data from subjects with brain disorders such as depression, Alzheimer’s disease and schizophrenia and to control subjects who have been matched using behavioural data. By comparing change points and partition specific networks, the *NEDM* may lead to the robust identification of cognitive states at rest for both controls and subjects with these disorders. It is hoped that the large-scale temporal features resulting from the accurate description of brain connectivity from our novel method, which might lead to better diagnostic and prognostic indicators of the brain disorders. More specifically, by comparing the change points of healthy controls to patients with these disorders, we may be able understand the key differences in functional brain processes that may eventually lead to the identification of biomarkers for the disease.

As an extension to monitoring change points in graph objects, we intend to incorporate higher order structures, such as tensors, in the network snapshot characterization procedure of the *NEDM*. Moreover, we intend to extend our analysis to include the estimation of network summaries that are based on the *local* topology and geometry of the graph. In particular, we intend to incorporate a motif-based analysis (Milo et al., 2002; Dey et al., 2019; Sarkar et al., 2019) and the concepts of topological data analysis (TDA), particularly, persistent homology, in the derivation of graph summary statistics (Carlsson, 2009; Patania et al., 2017; Carlsson, 2019). Indeed, tracking local network topological summaries based on graph persistent homology offers multi-fold benefits. First, this approach enables us to consider edge-weighted networks. Second, it allows for enhancing analysis of the underlying network organization at multi-resolution levels. Third, simultaneously considering multiple local network topological statistics based on graph persistent homology minimizes the loss of network information that currently occurs due to reducing a high dimensional structure to a univariate time series representation of a single network summary.

Another interesting theoretical direction is to explore various types of regularized approximation models (Gel and Barabanov, 2007; Bickel and Gel, 2011; Politis, 2015) for dynamics of local and global network topological summaries, and the associated error bounds, which as a result, can also yield an insight on theoretical guarantees of resampling and subsampling procedures (Kreiss et al., 2011; Fragkeskou and Paparoditis, 2018) in application to (non)linear processes of network topological descriptors and related uncertainty quantification in network anomaly detection.

## SUPPLEMENTARY MATERIAL

### R Code and Data

The supplemental files for this article include files containing R code and data for reproducing all the simulated and empirical studies in the paper.

### Appendix

The supplemental files include an Appendix which contains the following: (i) Theorem 1 from Gombay (2008) and Notations table. (ii) Simulation analysis for the *NEDM* size and power with *P* = 50. (iii) Additional results for the *NEDM* applied to the Anxiety fMRI data under the remaining network summaries. (iv) Plot for the *KVFM* Enron email network analysis. (v) Analysis and discussion of case study (resting state fMRI data). (vi) Proofs for Theorem 1 and Corollary 1. (vii) Procedure for comparing asymptotic and sieve bootstrap distributions.

## Acknowledgements

The work of D.O.B and Y.R.G. was partially supported by the National Science Foundation (NSF) of the United States (grant numbers IIS 1633331, DMS 1925346 and DMS 1736368). The work of I.C was partially supported by Natural Sciences and Engineering Research Council of Canada (NSERC: Grant/Award Number: RGPIN-2018-06638) and the Xerox Faculty Fellowship, Alberta School of Business. The authors would like to thank Gagan S. Wig for stimulating discussions on brain networks.

## Supplement

## 1 Appendix A: Notation and Theorem

### Theorem 1

(Gombay (2008)) *Assume the sequence of observations* {*Y_t_*} *satisfy equation (1.1) with Gaussian i.i.d. white noise* {*ε_t_*}, *var*(*ε_t_*) = *σ*^2^, *and characteristic polynomial* 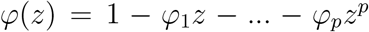 *has roots outside the unit circle. Then there exists a* (*p* + 2) - *dimensional Gaussian process* **B**(*u*) *with independent Brownian bridge components B*^(*r*)^(*u*), *j* = 1,…, *p* + 2, *such that*:

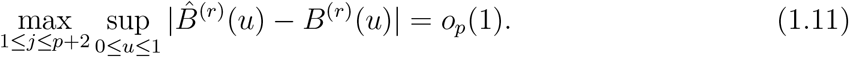

## 2 Appendix B: Experimental data analysis

### 2.1 Simulation study: Size & Power of the *NEDM* with *P* = 50

We performed Monte Carlo simulations (5000 iterations) under a no-change point (change in mean) scenario and a one-change point (change in variance) scenario with *P* = 50 using the same parameters stated in Pages 13-14 described under Section 4 of the manuscript. The results are presented in Figures 1 and 2, respectively.

**Figure 1:**
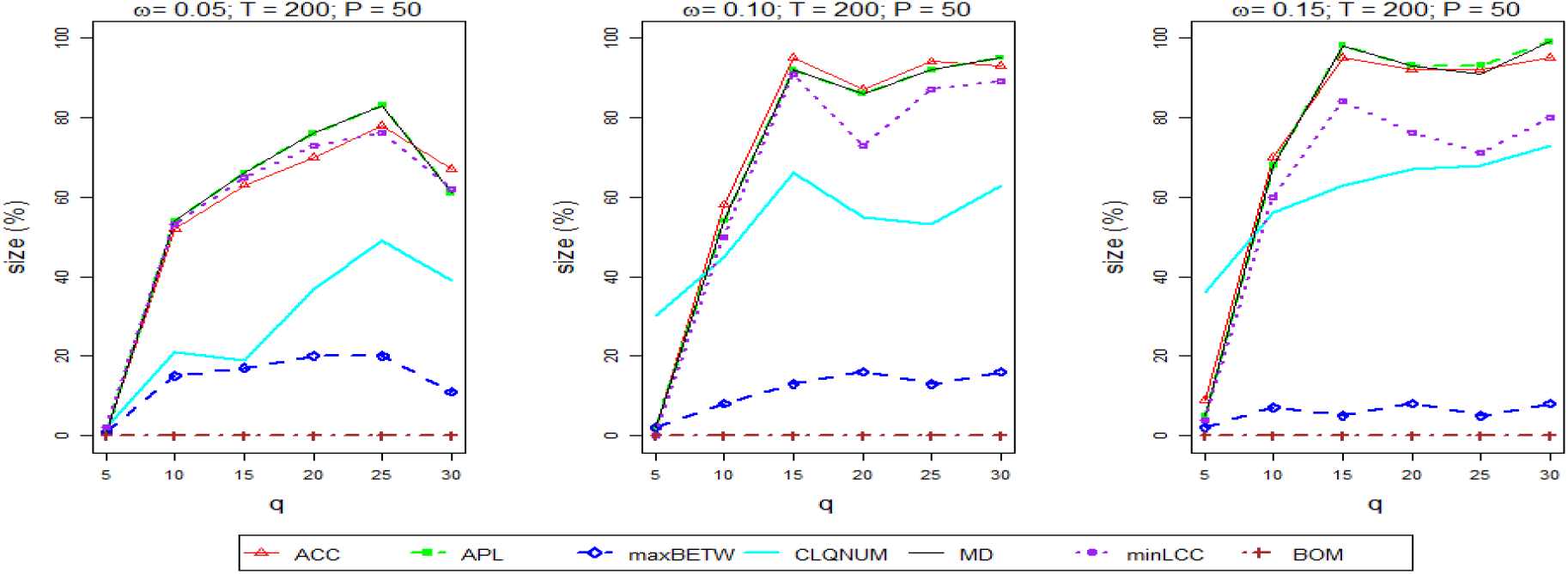
The size of the test for the no-change (change in mean) scenario with *T* = 200, *ω* = {0.05, 0.1, 0.15}, *q* = {5, 10, 15, 20, 25, 30} and *P* = 50.

In Figure 1, we find that as the window length (*q*) and the threshold *ω* increase the size of the test reported by all network summary statistics (with the exception of *maxBETW*) under *NEDM* also increase, leading to more liberal results. With *maxBETW*, however, we find that as both *q* and *ω* increase, the size of the test steadily approaches the declared 5% level of significance. In turn, *BOM* appears to be very conservative, delivering a size of the test close to 0% for all *q* and *ω*. This is in contrast to our analysis in the manuscript with *P* = 5. Here we found that the size of the test under *BOM* tends to be more liberal (ranging between 18% to 20%).

**Figure 2:**
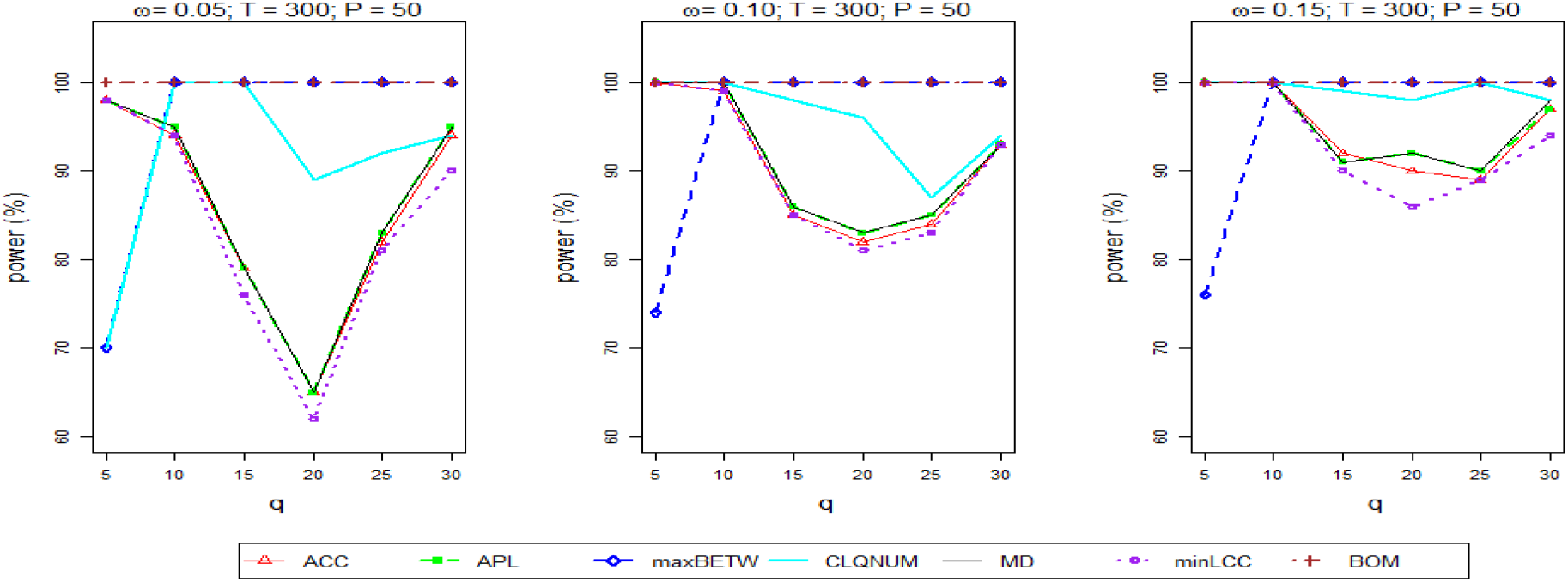
The power of the test for the one-change (change in variance) scenario with *T* = 300, *ω* = {0.05, 0.1, 0.15}, *q* = {5,10,15, 20, 25, 30}, *τ* = 157 and *P* = 50.

Figure 2 shows that, for *P* = 50, *NEDM* with the *maxBETW* network summary performs as well as *BOM* in terms of power (*i.e*., it correctly flags the one-change scenario every time) except for one combination. This is very similar to the results for *P* = 5. In summary, we can conclude that both the power and the size of the test tend to be sensitive to the choice of parameters (*q*, *ω* and *T*) and that the performance of the *NEDM* also depends on the choice of network summary statistic which in turn is likely to be driven by the underlying network topology.

### 2.2 Case study: Anxiety fMRI data

Figure 3 presents additional results for the Anxiety fMRI data based on the *NEDM* with *CLQNUM* and MD as the network summary statistics. To this end we find that the *NEDM* is able to detect 3 change locations with the *MD* summary, and 1 change location with the *CLQNUM* network summary. Table 2 presents a summary of the numerical results obtained based on this data.

**Table 2:** Summary of change points detected for the Anxiety fMRI data.

| Subject | *ACC* | *APL* | *maxBETW* | *CLQNUM* | *MD* | *minLCC* | *BOM* | *KVFM* |
| --- | --- | --- | --- | --- | --- | --- | --- | --- |
| 1 | 94 | 62,93 |  |  |  | 94 |  | 92 |
| 2 |  |  |  |  |  | 72 |  | 111,112,120 |
| 3 | 138 | 134 | 134 | 72 | 134 | 134 |  | 92,94 |
| 4 |  |  |  |  | 115 |  |  | 69,70,76,117 |
| 5 | 63 | 63,70 | 68,131 | 63 | 64 | 64 |  | 108 |
| 6 |  |  | 106 |  |  |  |  | 90 |
| 7 | 63 |  |  |  |  |  | 61 |  |
| 8 |  | 109 |  |  |  |  | 102 | 77,78 |
| 10 | 99 | 126 |  |  |  |  | 67 |  |
| 11 |  |  |  |  |  |  |  | 92,93 |
| 12 |  | 99 |  |  |  |  | 116 | 75,90,94 |
| 13 |  | 65,66 |  |  | 71 |  |  | 62,65 |
| 14 | 106 | 104 |  | 97 |  |  |  | 88,111 |
| 15 |  |  |  |  |  |  |  | 65,66,98 |
| 16 | 140 |  |  |  |  |  | 92 |  |
| 17 |  | 77 | 97 |  |  | 97,104 |  |  |
| 18 |  |  |  |  | 130 |  |  |  |
| 19 |  |  | 93 |  |  |  |  |  |
| 20 | 65 |  |  |  | 70 |  | 100 | 61,62,86 |

\* Subjects not included have no change points detected across all methods.

#### Results for Anxiety fMRI data

**Figure 3:**
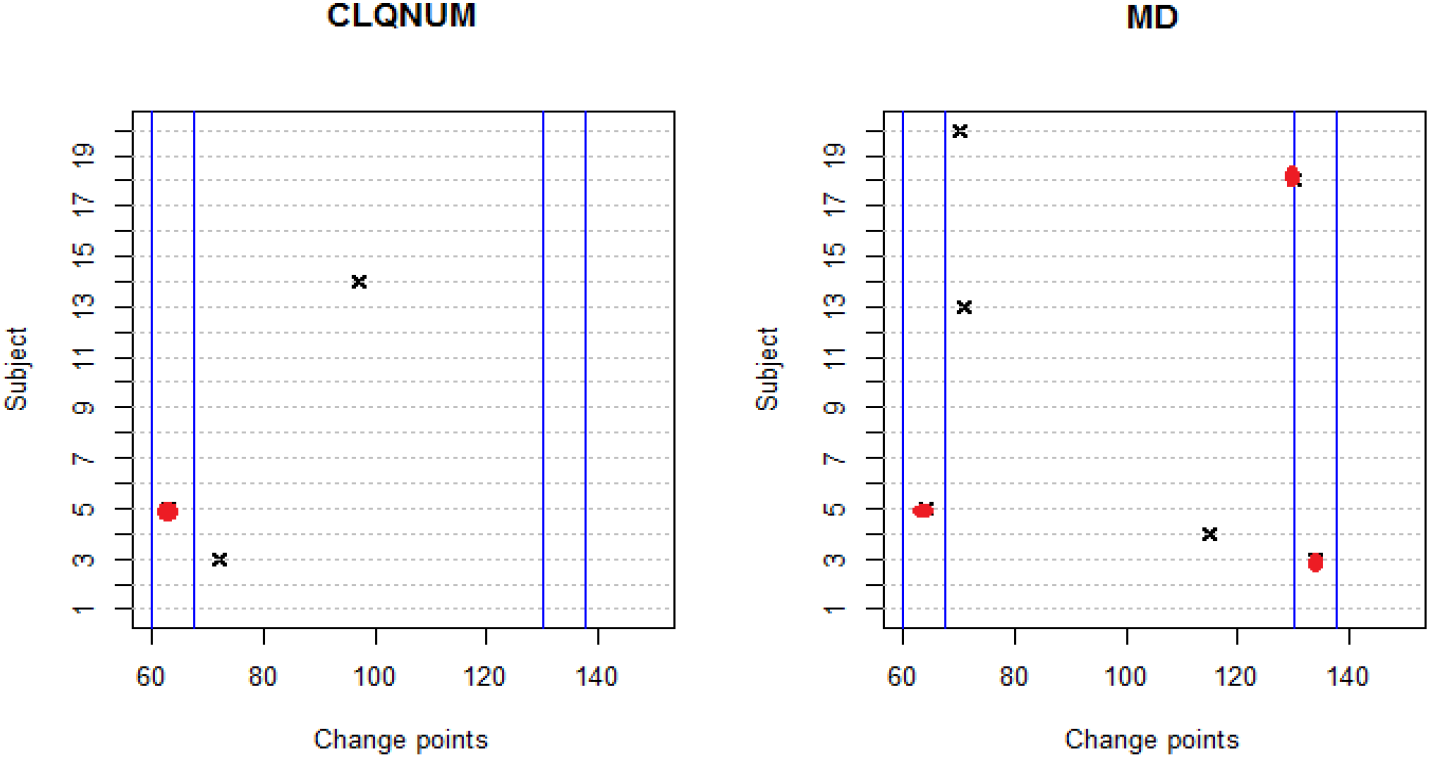
The change points detected for the Anxiety fMRI data using the *NEDM* and the *CLQNUM* and the *MD* summary statistics.

### 2.3 Case study: Enron email network using the *KVFM*

**Figure 4:**
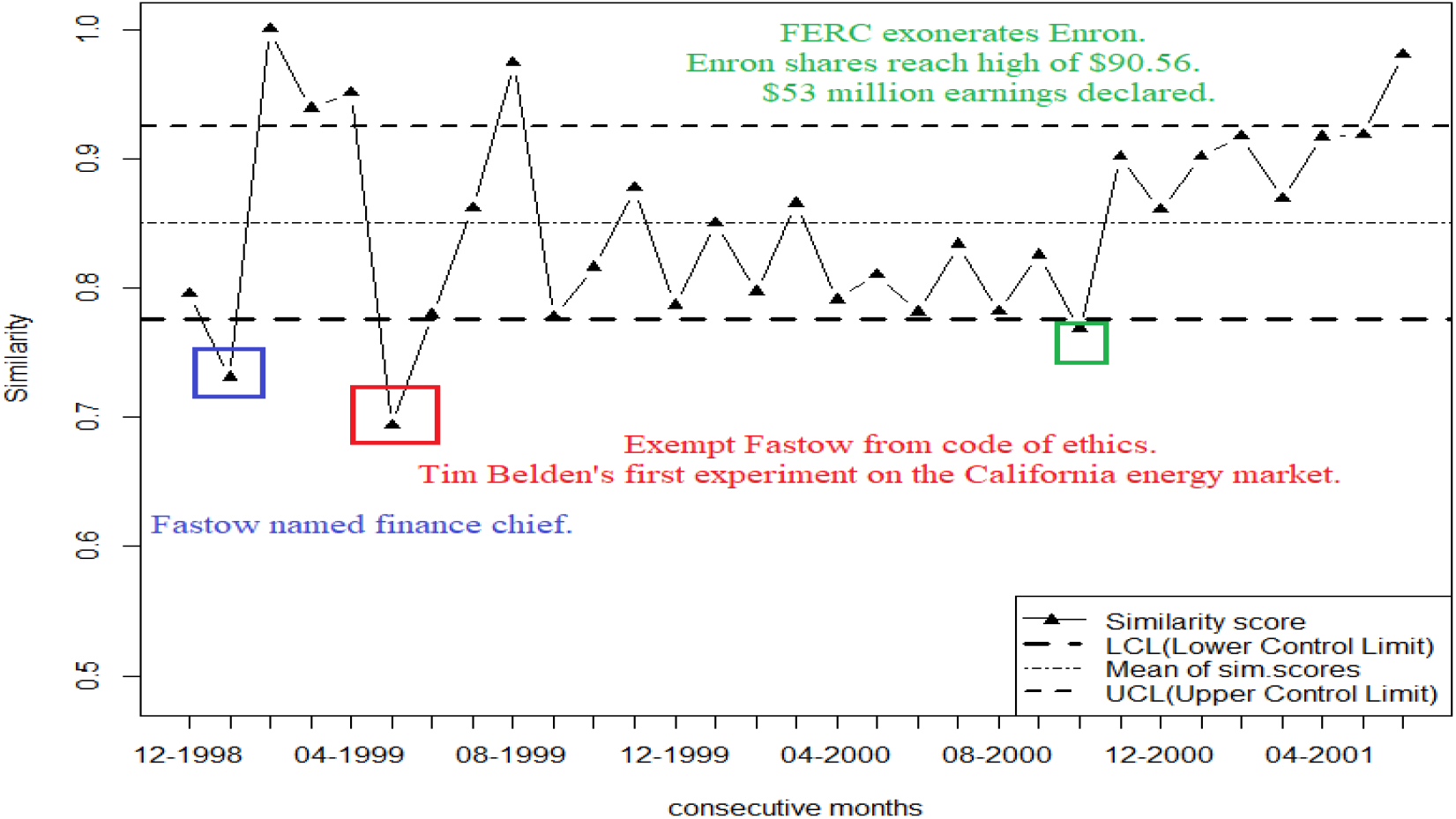
The detected change points for the Enron email data set from November 1998 – July 2001 using the *KVFM*.

### 2.4 Case study: resting state fMRI data

We apply the *NEDM*, with all network summary statistics except the *CLQNUM*, to a resting-state fMRI data as described in Cribben and Yu (2017). The *CLQNUM* summary is omitted due to computational challenges encountered during the analysis. Participants (*n* = 45) are instructed to rest in the scanner for 9.5 minutes, with the instruction to keep their eyes open for the duration of the scan. We apply the Anatomical Automatic Labeling (Tzourio-Mazoyer et al., 2002) atlas to the adjusted voxel-wise time series and produce time series for 116 Regions of Interest (ROIs) for each subject by averaging the voxel time series within the ROIs. In total, each time series contained 285 time points (9.5 minutes with *TR* = 2 *s*).

#### Results

Results obtained in this case study (as seen in Figure 5) show significant change point locations for all 45 subjects. As resting state data is unconstrained, we do not know where the true network change points occur. However, the *NEDM* finds significant change points in the network structure for many subjects with the maximum number of change points being 4 which lines up with the results in Cribben and Yu (2017). From Figure 5, we find a high indication that change points vary highly across network summaries and that some change states endure longer periods while others transition more quickly. On the other hand, results by the *BOM* and by the *KVFM* appear to suggest the detection of a much larger number of significant change points for each subject. This is expected for the *KVFM* given that it does not provide a quanitifier for statistical significance. Furthermore, judging from the close proximity and the total number of change points reported by these two methodologies, we deduce that some of the change points for the subjects are unrealistic for resting state fMRI data. As noted above, having change points so close to one another makes it impossible to estimate the network structure between each pair of significant change points.

**Figure 5:**
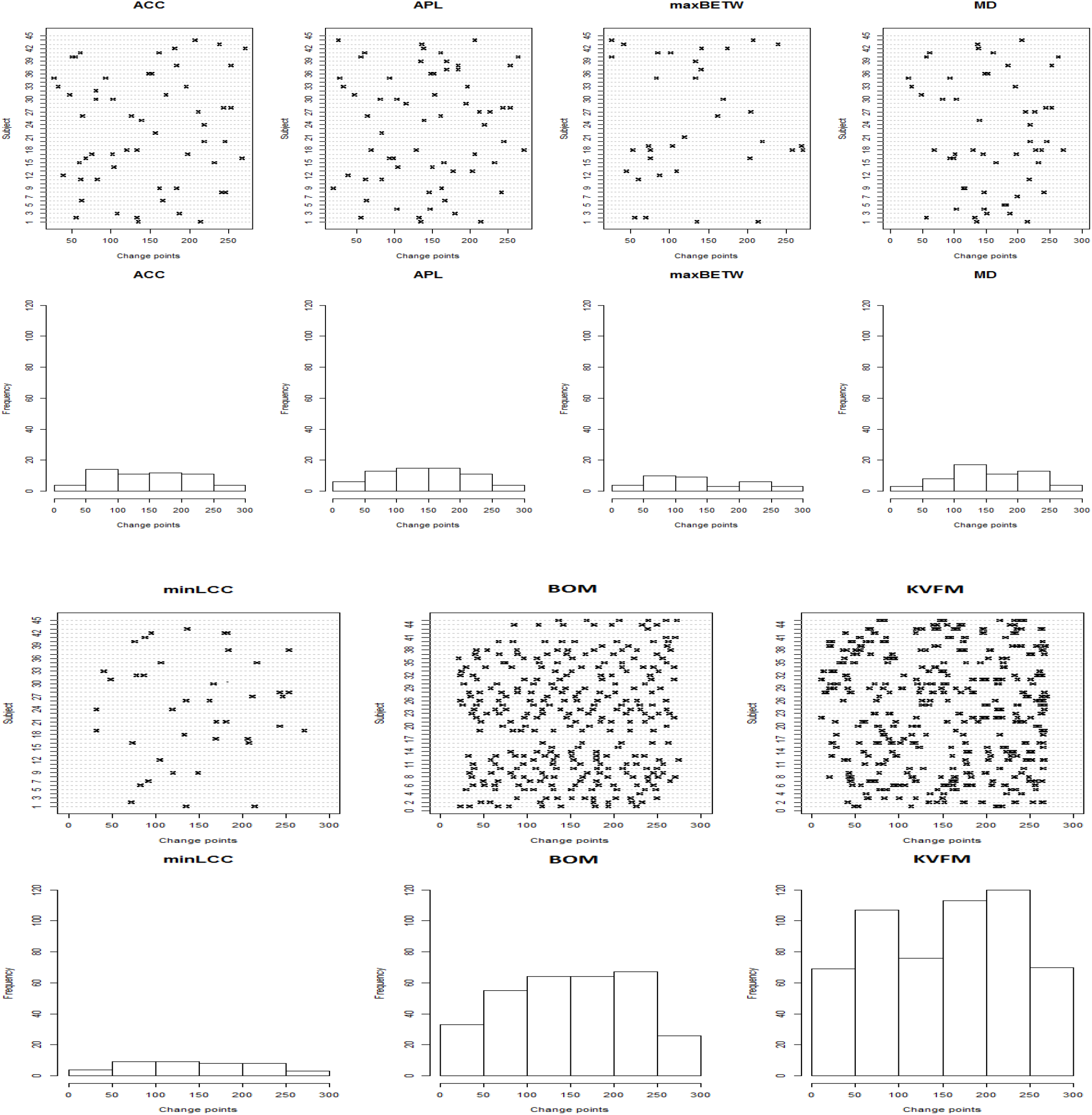
The detected change points for the resting state fMRI data set.

## 3 Appendix C: Proofs for Theorem 1 and Corollary 1

### Theorem 1

*Proof*. Note that the assumptions on *Y_t_* imply that it satisfies assumptions *A*1′, *A*2, and *B* of Bühlmann et al. (1997).

From Equation (1.2), we know that

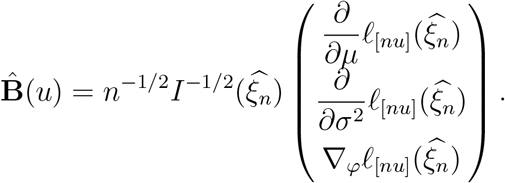

Hence, each individual component of *B*^(*r*)^(*u*) and its bootstrap counterpart *B*^(*r*)*^(*u*) also satisfy the assumption *C* of Bühlmann et al. (1997). As a result, by invoking Theorem 3.3 of Bühlmann et al. (1997) (as *n* → ∞), we obtain

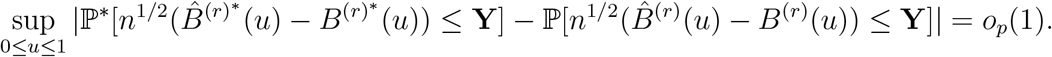

### Corollary 1

*Proof*. Note that 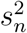 is a consistent estimator of 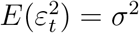 (Lehmann and Casella, 2006).

Hence, given that 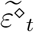 are independent random variables drawn from 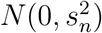,

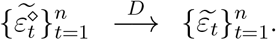

As a result,

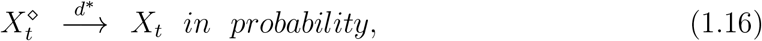

and in view of the proof for Theorem 1

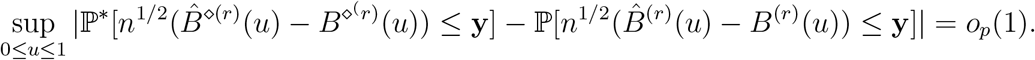

## 4 Appendix D: Asymptotic and Sieve bootstrap distribution simulations

The procedure for assessing the performance of the sieve bootstrap technique to the asymptotic distribution is set up as follows:

1. Simulate 5, 000 Monte Carlo simulations of an *AR*(1) time series with different samples (*n* = 100, 200, 300) and change point times (*τ* = 50, 100, 180).
2. Consider the following scenario for identifying change points in the *AR*(1) time series: change in mean (*μ*). We use Γ = 5000 replications for the sieve bootstrap procedure (as expressed in Algorithm 1).
3. With equation (1.1) as reference, the *AR*(1) model is defined as

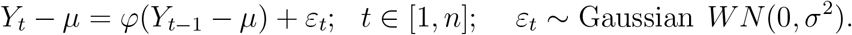
4. For each sample *n*, with *σ*^2^ and *φ* as nuisance parameters, we test the following:

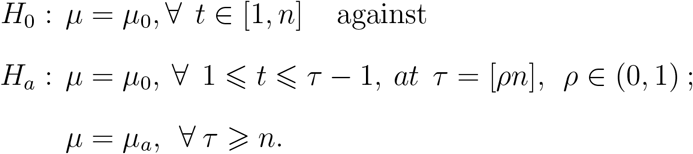
5. With *σ*^2^ = 1 and *μ*_0_ = 0, we use the values *φ* = (−0.5, 0.1, 0.5).

## Footnotes

1 In the experimental analysis we select parameters *q* and *ω* based on previous neuroscience studies. Alternatively, *q* and *ω* can be selected via cross-validation.

2 From Gombay (2008), a specific value of *φ* (and not |*φ*| or *φ*^2^) generates a specific change point statistic 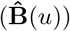 and the change point estimator 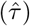.

